# Contributions of the N-terminal intrinsically disordered region of the SARS-CoV-2 nucleocapsid protein to RNA-induced phase separation

**DOI:** 10.1101/2022.03.30.486418

**Authors:** Milan Zachrdla, Adriana Savastano, Alain Ibáñez de Opakua, Maria-Sol Cima-Omori, Markus Zweckstetter

## Abstract

SARS-CoV-2 nucleocapsid protein is an essential structural component of mature virions, encapsulating the genomic RNA and modulating RNA transcription and replication. Several of its activities might be associated with the protein’s ability to undergo liquid-liquid phase separation. N^SARS-CoV-2^ contains an intrinsically disordered region at its N-terminus (NTE) that can be phosphorylated and is affected by disease-relevant mutations. Here we show that NTE deletion decreases the range of RNA concentrations that can induce phase separation of N^SARS-CoV-2^. In addition, deletion of the prion-like NTE allows N^SARS-CoV-2^ droplets to retain their liquid-like nature during incubation. We further demonstrate that RNA-binding engages multiple parts of the NTE and changes NTE’s structural properties. The results form the foundation to characterize the impact of N-terminal mutations and post-translational modifications on the molecular properties of the SARS-CoV-2 nucleocapsid protein.

**Statement:** The nucleocapsid protein of SARS-CoV-2 plays an important role in both genome packaging and viral replication upon host infection. Replication has been associated with RNA-induced liquid-liquid phase separation of the nucleocapsid protein. We present insights into the role of the N-terminal part of the nucleocapsid protein in the protein’s RNA-mediated liquid-liquid phase separation.

## Introduction

The severe acute respiratory syndrome coronavirus 2 (SARS-CoV-2) has caused an outbreak of a pandemic. The SARS-CoV-2 lipid membrane envelopes a positive-sense, single-stranded 30kb large RNA genome, which is encapsulated by the nucleocapsid protein (N^SARS-CoV-2^).

N^SARS-CoV-2^ is highly abundant and in addition to its structural role, it might play an important role in SARS-CoV-2 replication via RNA-driven liquid-liquid phase separation (LLPS) ^1,2,3,4,5,6^. From a therapeutic view, targeting N^SARS-CoV-2^ could be useful to improve the efficacy of currently available vaccines ^7,8^.

N^SARS-CoV-2^ consists of two structured domains flanked by intrinsically disordered regions (Figure 1A). The structured C-terminal domain is responsible for N^SARS-CoV-2^ dimerization and oligomerization ^9^. The flexible linker between the two folded domains involves a serine-arginine-rich region (SR-region), which mediates interaction between the nucleocapsid protein and various protein and RNA partners ^6,10,11^. Due to the high flexibility of the SR-rich linker region, the N-terminal folded domains are largely decoupled from the C-terminal dimer ^2,12,13^, giving the N-terminal domains enough flexibility to engage in RNA binding. Key to RNA-binding is a finger-like basic patch in the N-terminal domain ^14^. In addition, essentially all parts of the nucleocapsid protein have been shown to bind RNA or be involved in binding other partners ^14^.

**Figure 1.**
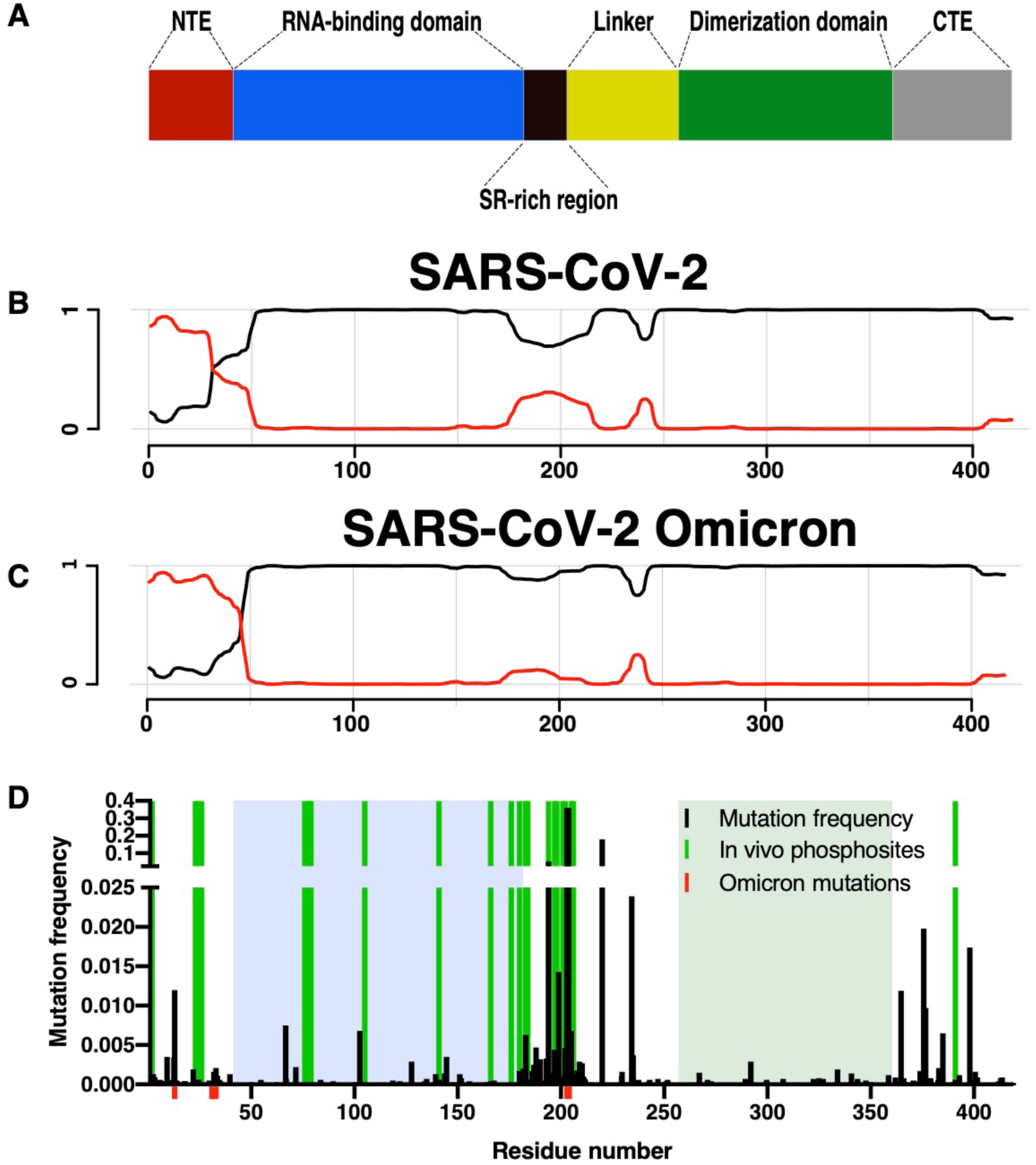
Sequence analysis of the nucleocapsid protein from the SARS-CoV-2 coronavirus. A) Domain organization of N^SARS-CoV-2^. B-C) Prediction of prion-like domains in the nucleocapsid protein of SARS-CoV-2 (B) and the Omicron variant (C) using the PLAAC algorithm ^19,20^. Background and prion-like domains are indicated by black and red lines, respectively. D) N^SARS-CoV-2^ mutation and phosphorylation analysis. Amino acid frequency of mutations using 219 909 sequences from the China National Center for Bioinformatics. Mutations present in the Omicron variant of SARS-CoV-2 are shown in red. Phosphorylation sites are highlighted in green. Folded domains are highlighted by blue and green boxes.

Most structural biology-focused research has so far concentrated on the two folded domains of N^SARS-CoV-2^, which are conserved between different coronaviruses ^15,16^. Less attention on the other hand has been paid to the disordered regions of N^SARS-CoV-2^, despite their suggested roles for modulating N^SARS-CoV-2^ activity. For example, the N-terminal intrinsically disordered tail (NTE) has been suggested to be responsible for the recruitment of N^SARS-CoV-2^ into stress granules ^17^. To close this gap, we characterize here NTE’s contribution to N^SARS-CoV-2^ liquid-liquid phase separation and binding to RNA, and provide insights into its molecular properties using a combination of phase separation assays, NMR spectroscopy and molecular dynamics (MD) simulations.

## Results

The ability of proteins to undergo liquid-liquid phase separation might be connected to sequence properties found in prion-like proteins ^18^. We applied the PLAAC ^19,20^ algorithm to identify amino acid sequences with prion-like composition within the N^SARS-CoV-2^ sequence. The analysis reveals pronounced prion-like properties in the disordered N-terminal tail and SR-linker region of N^SARS-CoV-2^ (Figure 1B), in agreement with previous analyses ^5,21^. Closer inspection shows that the N-terminal part of N^SARS-CoV-2^ contains several glutamine residues, known to favour prion-like sequence properties.

An important property of viruses is their rapid adaptation through mutations. To determine if the NTE of N^SARS-CoV-2^ not only has prion-like, potentially LLPS-promoting properties but also harbors mutations, we analyzed 219 909 N^SARS-CoV-2^ sequences. We find that most mutations are localized in the intrinsically disordered regions, particularly in the SR-rich linker region (Figure 1D). Mutations in the SR-region were suggested to increase the spread of the SARS-CoV-2 virus ^22^. Notably, the currently predominant SARS-CoV-2 variant B.1.1.529 (Omicron; Figure 1C) shares some mutations in the SR-region with previously characterized variants. In addition, the Omicron variant contains a P13L mutation and a Δ^31^ERS^33^ deletion which enhances the prion-like properties of the NTE (Figure 1D). Further support for an important role of the NTE in the life cycle of N^SARS-CoV-2^ comes from the observation that, besides the SR-region, it can be phosphorylated in SARS-CoV-2 infected cells (Figure 1D) ^[23,24]^.

Motivated by the above sequence analysis we investigated the regulatory effect of the NTE on the LLPS of N^SARS-CoV-2^. While some studies probed the importance of the NTE for LLPS and biomolecular condensation of N^SARS-CoV-25,17,25^, its precise role remains unclear. One study suggested that the NTE contributes to the recruitment – potentially through LLPS – into stress granules ^17^. Other studies, however, showed that multiple regions of N^SARS-CoV-2^ are important for RNA-induced LLPS ^5,25^. To gain additional insights into the contribution of the NTE to LLPS of N^SARS-CoV-2^, we recombinantly prepared two constructs: the 419-residue full-length N^SARS-CoV-2^ protein and a 379-residue construct with deleted NTE (ΔNTE^SARS-CoV-2^; Figure 2A). Deletion of the NTE changes the net charge of the protein at pH 7.5 from +25 for the full-length protein to +20 for ΔNTE^SARS-CoV-2^.

**Figure 2.**
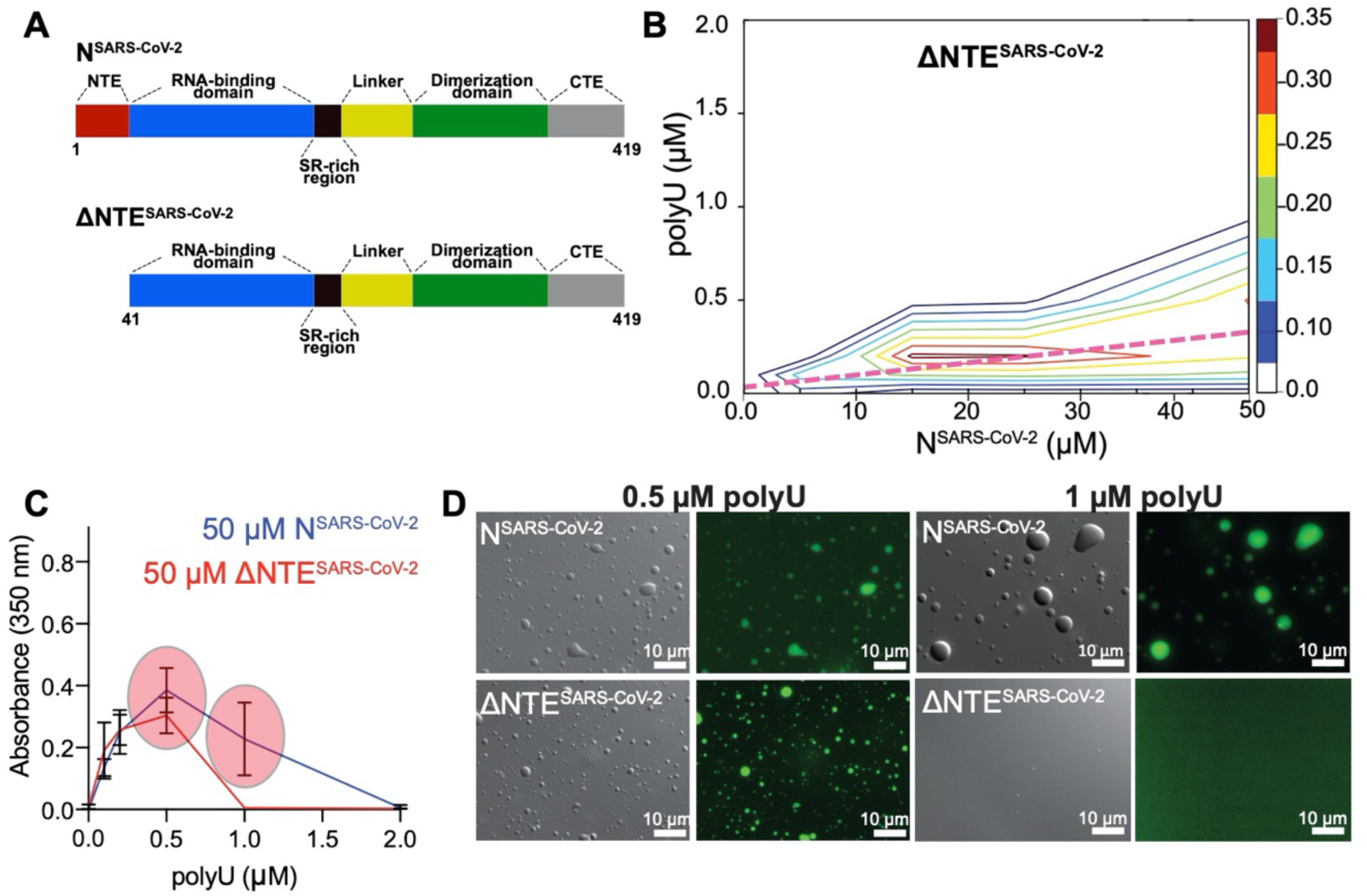
The N-terminal disordered region of the nucleocapsid protein of SARS-CoV-2 promotes liquid-liquid phase separation. A) Domain organization of full-length N^SARS-CoV-2^ and the N-terminally truncated ΔNTE^SARS-CoV-2^ protein. B) Phase diagram of ΔNTE^SARS-CoV-2^ at protein concentration up to 50 μM and in the presence of increasing polyU RNA concentrations measured as turbidity at 350 nm. Conditions of charge compensation are marked by the dashed magenta line. C) Absorbance at 350 nm of 50 μM full-length N^SARS-CoV-2^ or of 50 μM ΔNTE^SARS-CoV-2^ in the presence of 0.1, 0.2, 0.5, 1, 2 μM polyU RNA. Red circles highlight differences in the LLPS behaviour between the two constructs. D) DIC/fluorescence microscopy images displaying the differences in the phase separation behavior between full-length N^SARS-CoV-2^ and ΔNTE^SARS-CoV-2^.

Using these two constructs, we analyzed the effect of the NTE deletion on LLPS for different protein and RNA concentrations. We measured sample turbidities at 350 nm of both full-length N^SARS-CoV-2^ and ΔNTE^SARS-CoV-2^ at protein concentrations up to 50 μM, and in the presence of 0-2 μM polyU (800 kDa), which serves as a simple substitute for viral RNA (Figure 2B-C and Figure S1). The obtained phase diagram of the full-length protein is in agreement with our previously published data (Figure S1) ^6^. In the case of the truncated protein, we also observed maximum turbidity in conditions close to charge neutralization (Figure 2B). However, when we compare in detail the dependence of turbidity on RNA concentration for 50 μM of protein, a clear difference in LLPS behavior is observed between the full-length and the N-terminally truncated protein: only the full-length protein undergoes LLPS in the presence of 1 µM polyU (Figure 2C). In agreement with the turbidity measurements, abundant protein-dense droplets were seen by differential interference contrast (DIC) and fluorescence microscopy for both proteins at 0.5 µM polyU, but only for the full-length protein at 1 µM polyU (Figure 2D).

We previously reported that N^SARS-CoV-2^ droplets become less dynamic over time ^6^. We therefore repeated the analysis with ΔNTE^SARS-CoV-2^ by measuring fluorescence recovery after photobleaching (FRAP) with freshly prepared droplets in presence of charge-compensating amounts of polyU, followed by a second measurement after one hour waiting time (Figure 3). In contrast to the full-length protein, we observed similar profiles for early and late time points of ΔNTE^SARS-CoV-2^ droplets, i.e. that no change of dynamics over time occured. Deletion of the NTE thus allows N^SARS-CoV-2^ droplets to retain their liquid-like nature.

**Figure 3.**
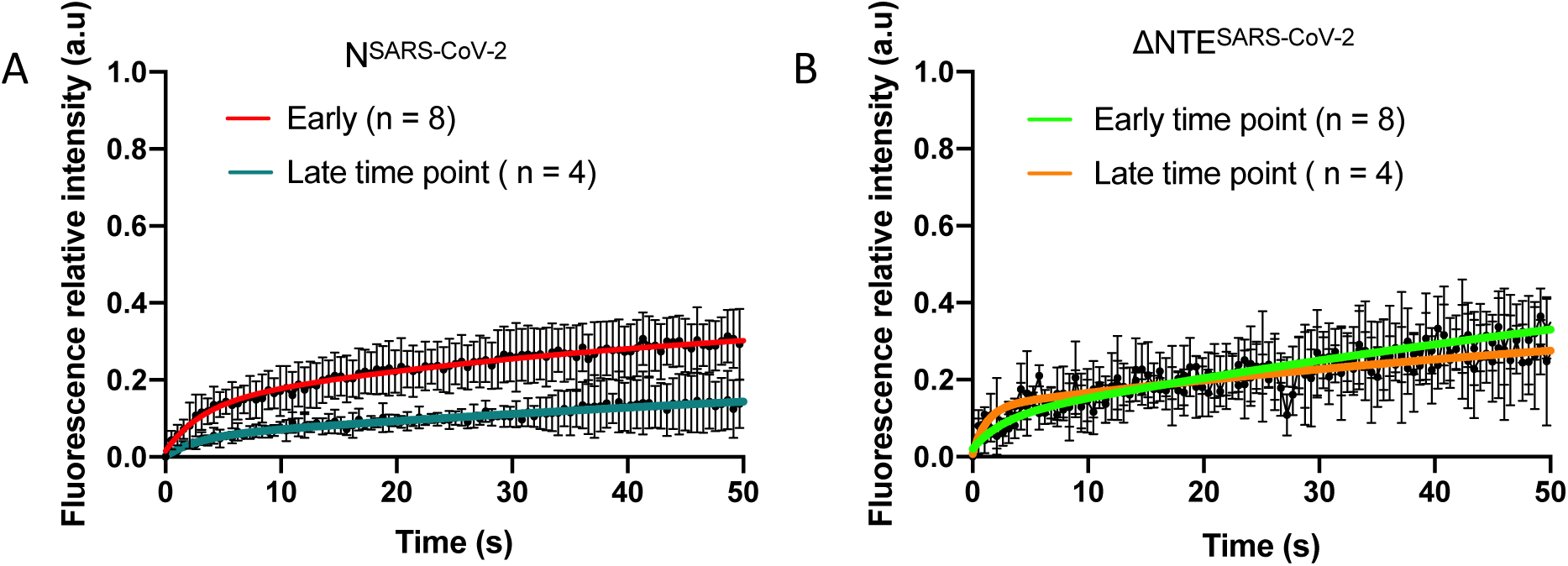
Time-dependent change in diffusion of phase-separating N^SARS-CoV-2^. A) Fluorescence recovery of 30 µM N^SARS-CoV-2^ droplets formed in presence of 0.3 µM polyU measured at early (red) and late (blue) time points. Error bars represent standard deviations for n = 8 and n= 4 for early and late time points, respectively. Scale bars, 10 µm. B) Fluorescence recovery of 30 µM ΔNTE^SARS-CoV-2^ droplets formed in presence of 0.3 µM polyU measured at early (green) and late (orange) time points. Error bars represent standard deviations for n = 8 curves and n= 4 for early and late time points, respectively.

To better understand the molecular basis of the influence of the NTE on the RNA-induced phase separation of N^SARS-CoV-2^, we characterized the structural properties of the NTE using NMR spectroscopy and molecular dynamics simulations (MD). Based on previous studies, already demonstrating that the NTE is disordered ^5,26,27^, we prepared a synthetic peptide that comprises the NTE of N^SARS-CoV-2^. Focusing on the isolated NTE region has the advantage of decreased NMR signal overlap, which arises from the protein’s intrinsically disordered linker and C-terminal regions (Figure 4A).

**Figure 4.**
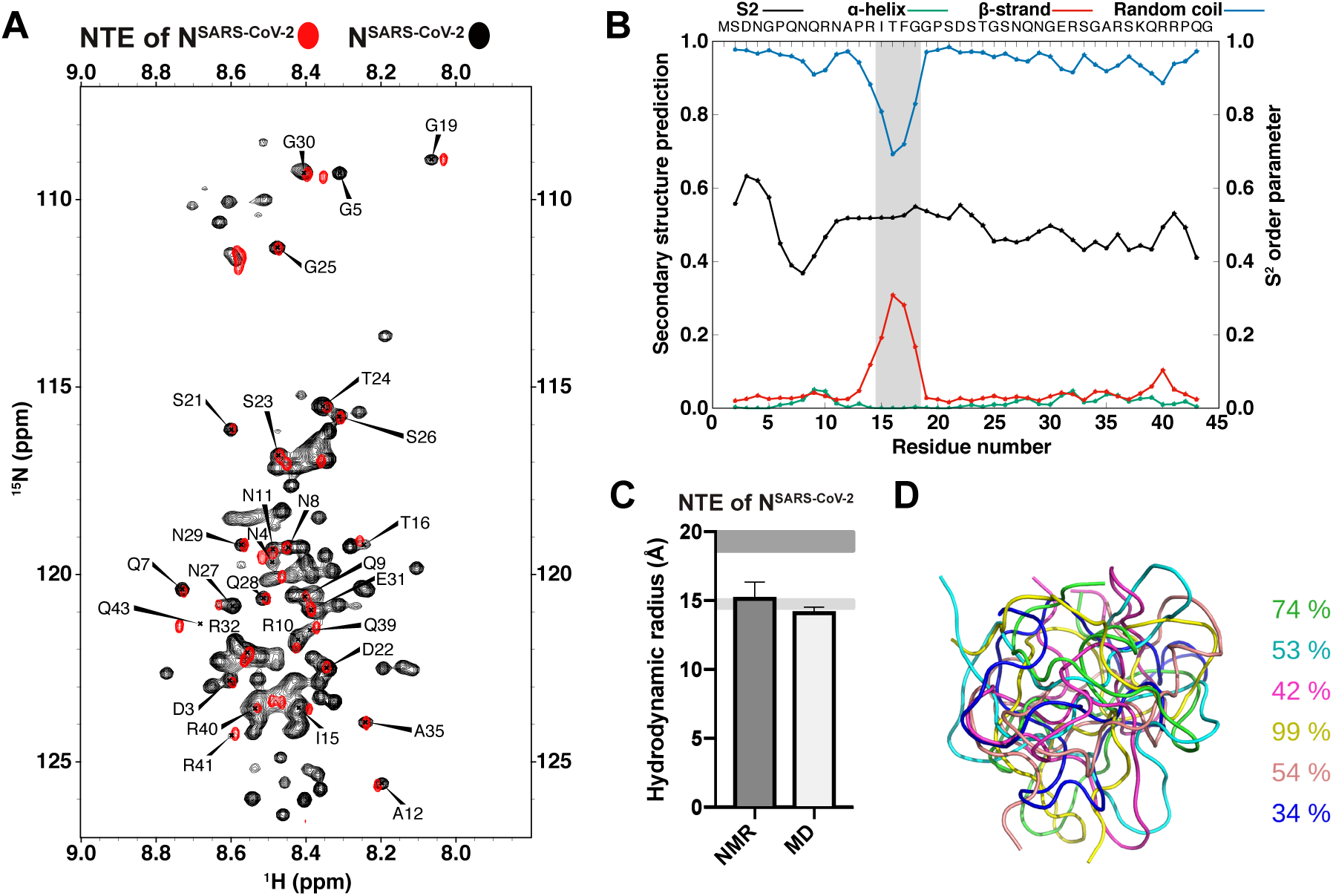
The N-terminal region of the nucleocapsid protein of SARS-CoV-2 forms a compact structure with transient β-structure. A) Superposition of ^1^H-^15^N HSQC spectra of full-length N^SARS-CoV-2^ in yellow and the NTE of N^SARS-CoV-2^ in red measured at 5 °C with amino-acid assignments. B) Order parameter, S^2^, and secondary structure prediction from the experimental chemical shifts of the NTE peptide of N^SARS-CoV-2^ using the TALOS+ software ^28^. The region with transient β-structure is highlighted in grey. C) Hydrodynamic radius of the NTE of N^SARS-CoV-2^ determined by NMR and MD simulations. The upper (dark grey) and lower (light grey) horizonal bars indicate the estimated ranges for a denatured and a globular protein, respectively ^29^. D) Representative structures of the NTE of N^SARS-CoV-2^ obtained from six independent MD simulations. The occupancy of the most populated cluster in each of the six MD simulations is indicated on the side.

Figure 4A shows a superposition of the natural abundance ^1^H-^15^N spectrum of the NTE peptide with that of full-length N^SARS-CoV-2^ at 5 °C. At this low temperature, only residues from the disordered regions of N^SARS-CoV-2^ are observable, while the folded N- and C-terminal domains are broadened beyond detection (Figure 4A). Overall more cross-peaks are seen in the spectrum of full-length N^SARS-CoV-2^, in agreement with extensive dynamics in the NTE, the SR-rich linker region and the C-terminal tail. In addition, good agreement is present between many of the cross-peaks of the isolated NTE (red cross-peaks) and the corresponding cross-peaks of the ^1^H-^15^N spectrum of full-length N^SARS-CoV-2^ (black cross-peaks in Figure 4A).

We then obtained the sequence-specific assignment of the NTE peptide using a combination of two-dimensional NOESY and TOCSY spectra and mapped it onto the ^1^H-^15^N HSQC spectrum (Figure 4A). The analysis showed that deviations in peak positions between the isolated NTE and the full-length protein occur mainly at NTE’s termini. Notably, there is little signal overlap between the chemical shifts of a peptide corresponding to the SR-rich linker region and the cross-peaks in the ^1^H-^15^N HSQC of full-length N^SARS-CoV-2^ (Figure S2). In contrast to the NTE region, the SR-rich linker region likely has different conformational properties in the isolated state or interacts with one of the folded domains.

Next, we analyzed the presence of transient secondary structure in the NTE using the assigned NMR chemical shifts (Figure 4B). We find that residues 13-19 display a propensity to transiently populate extended/β-structure-like conformations. The propensity of residues 13-19 for extended/β-structure-like conformations is in agreement with chemical shifts reported for N^SARS-CoV-2^ constructs comprising the NTE as well as the N-terminal folded domain and the linker region ^26,27^, while a previously reported molecular dynamics (MD) simulation suggested the presence of a transient α-helix between residues 31-38 but did not identify any β-structure in the NTE ^2^. The secondary structure analysis based on chemical shifts did not provide any evidence for α-helical structure between residues 31-38 suggesting that its population might be very small or require additional binding partners for stabilization. In addition, we observed a pronounced compaction of the NTE in the MD simulation (Figure 4C). The hydrodynamic radius of the isolated NTE corresponds to the value expected for a globular structure of the same sequence length, in good agreement with experimentally determined hydrodynamic radius values (Figure 4C).

The above described influence of the NTE on RNA-induced phase separation of N^SARS-CoV-2^ (Figure 2), suggests that the NTE directly binds to RNA. To gain insight into a potential direct interaction between the NTE and RNA, we performed an NMR titration of full-length N^SARS-CoV-2^ with increasing concentrations of polyU. In agreement with the previously determined LLPS properties ^1,3,5,6^, the NMR sample became highly turbid when we added 0.5 μM polyU to 50 μM N^SARS-CoV-2^, and upon further increase to 1.5 μM polyU the sample turned transparent again. In parallel to the changes in sample turbidity, many cross-peaks disappeared from the ^1^H-^15^N HSQC of N^SARS-CoV-2^ (Figure S3), but partially reappeared at high RNA concentration (Figure 5A).

**Figure 5.**
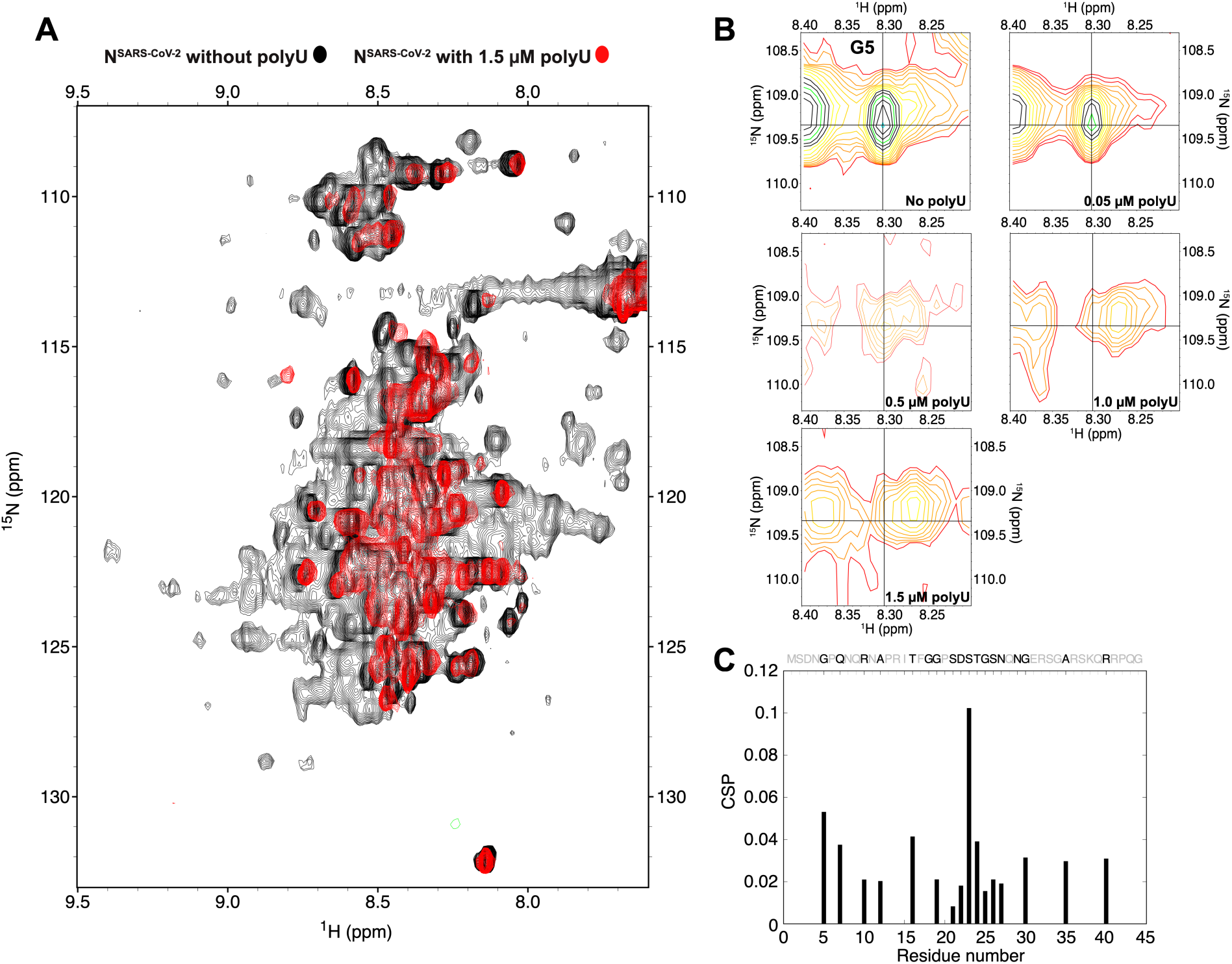
RNA-interaction of the N-terminal IDR region of N^SARS-CoV-2^. A) Superposition of ^1^H-^15^N HSQC spectra of full-length N^SARS-CoV-2^ without polyU RNA and in the presence of 1.5 μM polyU. Spectra were measured at 5 °C. B) ^1^H-^15^N HSQC spectra of the N^SARS-CoV-2^-residue G5 in the presence of 0, 0.05, 0.5, 1.0 and 1.5 μM polyU RNA. C) Chemical shift perturbation of the NTE upon interaction with 1.5 μM polyU RNA (NTE sequence shown above; unassigned residues are displayed in light grey).

The disappearance of the NMR signals at conditions of LLPS likely arise from increased relaxation rates in the protein when it is concentrated into the viscous phase of N^SARS-CoV-2^/RNA droplets ^30^. Additional ^1^H-^13^C XL-ALSOFAST HSQC experiments further showed that even for the more flexible methyl groups, RNA-induced LLPS of N^SARS-CoV-2^ causes severe line broadening (Figure S4).

Despite the strong LLPS-induced broadening of the cross-peaks in the ^1^H-^15^N HSQC spectra of N^SARS-CoV-2^, we could follow a few selected residues at increasing polyU concentrations. This is illustrated for the cross-peak of G5 in Figure 5B. In the absence of polyU, a single peak is observed. The peak weakens upon addition of 0.05 μM polyU, i.e. at the onset of LLPS (Figure 2B,C). At the conditions of maximum LLPS (0.5 μM polyU, as assessed by turbidity; Figure 2B,C) the cross-peak appears as doublet (Figure 5B). With further increase of the polyU concentration, the G5 signal at the original position is further broadened such that only the newly generated cross-peak remains at the highest tested polyU concentration (1.5 μM; Figure 5B). The observations suggest that we detect both an RNA-free conformation and an RNA-bound conformation in the NTE. The two conformations are in slow exchange on the NMR chemical shift time scale. In addition, the RNA-bound conformation of G5 remains sufficiently dynamic in order to be observable in conditions of strong LLPS. Similar observations were made for other cross-peaks in the NTE (Figure S5).

Further support for a direct interaction between the NTE of N^SARS-CoV-2^ and RNA is accessible at a concentration of 1.5 μM polyU, i.e. the high RNA concentration at which no LLPS is present anymore (Figure 2B,C). While the folded N-terminal domain remains invisible in this condition, likely due to strongly increased relaxation rates associated to its RNA-binding, many cross-peaks of NTE residues are visible (Figure 5A). Careful inspection of selected NTE cross-peaks, which are not severely affected by signal overlap, reveals chemical shift perturbations across the NTE sequence, with S23 displaying the highest chemical shift perturbation (Figure 5C). Notably, S23 of N^SARS-CoV-2^ has been shown to be phosphorylated in SARS-CoV-2-infected cells ^23,24^ suggesting that phosphorylation of the NTE can regulate N^SARS-CoV-2^‘s RNA-induced condensation and stress granule association in vivo.

## Discussion

N^SARS-CoV-2^ undergoes liquid-liquid phase separation with RNA ^1,2,3,5,6^. The formation of phase-separated N^SARS-CoV-2^/RNA-dense condensates might influence viral replication ^6,31^, drive genome packaging ^3^, or modulate the stress-response ^2,32^. It is therefore important to gain insight into the molecular structural basis of RNA-induced N^SARS-CoV-2^ phase separation. In the current study we show that the N-terminal intrinsically disordered region (NTE) modulates phase separation of N^SARS-CoV-2^ and changes the droplet ageing properties through direct binding to RNA. We further provide evidence that RNA-binding engages multiple parts of the NTE and changes its structural properties.

The N protein is an essential component of positive-sense coronaviruses, several of which have been identified in humans. While N protein’s two folded domains are well conserved among coronaviruses, the intrinsically disordered regions have little sequence conservation. They vary in both amino acid composition and sequence length and thus their physico-chemical properties. The NTE of N^SARS-CoV-2^ is rich in asparagine and glutamine residues, a typical feature of prion-like proteins (Figure 1B). There is growing evidence that prion-like regions play an important role in the development of neurodegenerative diseases through favoring protein aggregation ^35^. Notably, the mutations of the Omicron variant increase the content of the prion-like character in the NTE. We further show that deletion of the NTE weakens the ability of N^SARS-CoV-2^ to form liquid-like droplets with polyU RNA (Figure 2). Rather than abolishing LLPS, however, ΔNTE^SARS-CoV-2^ binding saturates at lower RNA concentrations. This can be rationalized on the basis of a lower net charge upon deletion of the NTE and the importance of charge neutralization for RNA-induced LLPS of N^SARS-CoV-26^. In addition, we show that the droplets of the truncated N protein remain fluid in contrast to the full-length protein ^6^. Considering that LLPS of N^SARS-CoV-2^ with the 3’ and 5’ ends of the viral genome promotes nucleocapsid assembly ^3,36^ and that the NTE is important for nucleocapsid assembly ^37^, viral viability ^38^, as well as binding the membrane protein ^39^, our study suggests that the NTE can contribute to diverse activities of N^SARS-CoV-2^.

In contrast to the N-terminal globular domain of N^SARS-CoV-240^, the NTE does not contain a known RNA binding motive. We however detected a direct interaction between the NTE and RNA. Electrostatic interactions are likely to be the main driving force for this interaction, in particular with a positively-charged patch at the C-terminus of the NTE, which contains three arginine and one lysine residue (Figure 2B). Consistent with an involvement of this positively charged region in RNA-binding, the cross-peaks of some of these residues disappeared in the presence of RNA due to intermediate chemical exchange. However, the NMR signal perturbations were not restricted to the C-terminus of the NTE, but spread over the whole NTE (Figure 5C). In addition, we observed the presence of two distinct conformations for several residues likely corresponding to an RNA-free and RNA-bound form. This indicates that RNA binding to the NTE is not restricted to residues capable of electrostatic interactions, in agreement with a recently suggested RNA-binding mechanism ^41^. We further note that the most affected residue of the NTE, S23, is phosphorylated in vivo ^23,24^, suggesting that phosphorylation of the NTE might influence N^SARS-CoV-2^‘s ability to phase separate. Consistent with this hypothesis, we previously showed that phosphorylation of the SR-region of N^SARS-CoV-2^ by the SRSF protein kinase 1 (SRPK1) influences N^SARS-CoV-2^ phase separation ^6^. Modulation of phosphorylation pathways thus might influence N^SARS-CoV-2^ condensation in infected cells and serve as a useful entry point for therapeutic intervention against COVID-19 ^42^.

### Experimental Section

#### Protein preparation

DNA encoding for N^SARS-CoV-2^ (UniProtKB - P0DTC9 obtained from ThermoFisher Scientific, GeneArt) was cloned using BamHI and HindIII restriction enzymes into a pET28 vector, comprising a Z2 solubility tag and a N-terminal 6× HIS-Tag for protein purification. BL21(DE3) competent cells were transformed with the cloned plasmids, with kanamycin resistance for colony selection, (ThermoFisher Scientific, Invitrogen) and grown in LB until reaching OD_600_ of 0.7-0.8. Cell cultures were then induced with 0.5 mM IPTG and grown overnight at 37 °C. For 15N-labeled protein the cells were grown in LB until reaching an OD_600_ of 0.7-0.8, then centrifuged and washed with M9 salts. Cells were subsequently resuspended in minimal M9 medium with 2g/l of 15NH_4_Cl as a sole nitrogen source. After 1 hour the cell cultures were induced with 0.5 mM IPTG and grown overnight at 37 °C. Grown cell cultures were harvested by centrifugation and either used directly or stored at -80 °C. Full-length plasmid was used as a template for PCR to clone DNA of ΔNTE^SARS-CoV-2^ to a new plasmid, which was subsequently expressed using the same protocol as for the full-length protein.

The cell pellets were resuspended in lysis buffer (25 mM HEPES, 300 mM NaCl, 1 mM EDTA, pH 8.0) supplemented with CaCl_2_, MnCl_2_, MgCl_2_, lysozyme, DNAse I (SigmaAldrich, Roche, 04536282001), and cOmplete™, EDTA-free Protease Inhibitor Cocktail (SigmaAldrich, Roche, 5056489001). All purification steps were performed at 4 °C to avoid precipitation.

The cells were lysed using sonication and the protein solution was cleared from cell debris by ultracentrifugation at 75,000 g at 4 °C for 30 minutes (Optima XPN-80, Beckman Coulter).

The supernatant was filtered with 0.2 μm filter and diluted to reduce the NaCl concentration to 100 mM. The sample was then loaded onto HiTrap SP HP column (Cytiva) and the bound protein was eluted with 1 M NaCl. The flow-through was loaded onto the column several times in order to remove bound nucleic acid, as indicated by a ratio A_260_/A_280_ > 1, until there was no protein left. Collected fractions with A_260_/A_280_ > 1 were prone to aggregate, and were therefore kept separately from the fractions with A_260_/A_280_ < 1. Both samples were then diluted to 500 mM NaCl concentration and loaded onto HisTrap FF column (Cytiva). The bound protein was eluted with 500 mM imidazole. After overnight dialysis (25 mM TRIS, 170 mM NaCl, 5mM imidazole, pH 8.0) the His-Tag was cleaved off by a TEV protease (as confirmed by SDS PAGE gel) and samples were loaded again onto HisTrap FF column. At this point, the protein was still binding to the column. Reasoning that a cut protein bound to the column via electrostatic interactions with a small portion of an uncut protein, the cut protein was eluted with 300 mM NaCl. The bound uncut protein was subsequently eluted by 500 mM imidazole. The last purification step was done using size exclusion chromatography, HiLoad 26/600 Superdex 200pg (buffer composition: 25 mM TRIS, 300 mM NaCl, pH 8.0).

The A_260_/A_280_ ratio of 0.55 measured in the final fractions confirmed the absence of nucleic acid contamination. Protein purity was verified using SDS page gel. Samples were then concentrated and exchanged to either NMR buffer (50 mM sodium phosphate, 10 % D_2_O,0.01 % NaN_3_, pH 6.8) or buffer for phase separation experiments (20 mM sodium phosphate, pH 7.6).

### Turbidity measurements

Phase diagrams of full-length N^SARS-CoV-2^ and ΔNTE^SARS-CoV-2^ were determined using a NanoDrop spectrophotometer (ThermoFisher Scientific, Invitrogen). PolyU (in concentrations from 0 to 2 μM) was added right before the experiments, followed by thoroughly pipetting and measurement of turbidity at 350 nm UV-Vis. Averaged turbidity values and the error bars were derived from measurements of three independent, freshly prepared samples.

### DIC/fluorescence microscopy

Full-length N^SARS-CoV-2^ and ΔNTE^SARS-CoV-2^ proteins were labeled using Alexa-fluor 488™ (green) microscale protein labeling kits (ThermoFisher Scientific, Invitrogen). Small amounts (∼0.3 µl) of fluorescently-labeled N^SARS-COV-2^ were premixed with unlabeled N^SARS-CoV-2^ and diluted to 30-50 μM final concentration with 20 mM sodium phosphate buffer at pH 7.5. 0.3 μM or 0.5 μM polyU RNA was added to N^SARS-CoV-2^ or ΔNTE^SARS-CoV-2^, respectively, achieving approximately charge compensation. 5 μl of the sample were subsequently loaded onto a slide and covered with a 18 mm coverslip. DIC and fluorescent micrographs were acquired on a Leica DM6B microscope with a 63× objective (water immersion) and processed using Fiji software (NIH).

### Fluorescence recovery after photobleaching (FRAP) measurements

FRAP measurements have been performed on 30 µM full-length N ^SARS-CoV-2^ or ΔNTE^SARS-CoV-2^ in presence of 0.3 µM polyU in 20 mM sodium phosphate buffer (NaP), pH 7.5 on a Leica TCS SP8 confocal microscope using a 63x objective (oil immersion) and a 488 argon laser line. Prior measurement, unlabeled FL-or 41-419 N^SARS-CoV-2^ were mixed with Alexa Fluor 488 lysine labeled FL-or 41-419 N^SARS-CoV-2^.

Fluorescence recoveries have been recorded as previously described ^1^. Upon addition of polyU, i.e. on freshly formed droplets early time points were recorded within a time <15 min, late time points have been acquired after one hour of sample incubation on the glass.

Regions of interest (ROIs) of ∼4 µm in diameter have been bleached with 50 % of laser power, recovery was imaged at low laser intensity (5%). 100 frames were recorded with one frame per 523 ms. The data have been processed using FIJI software (NIH).

FRAP recovery curves were obtained using standard protocols. Briefly, for each FRAP measurement intensities for pre-bleaching, bleaching and post-bleaching ROIs have been measured. A pre-bleaching ROI corresponded to a selected region in the droplet before bleaching; the bleached ROI corresponded to the bleached area while the reference ROI corresponded to an area which did not experience bleaching. The fluorescence intensity measured for each of the described ROIs was corrected by background subtraction: a region where no fluorescence was detected was used to calculate the background. Thus, the FRAP recovery was calculated as:

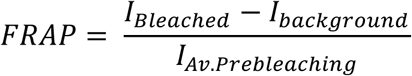

The value obtained was then corrected by multiplication with the acquisition bleaching correction factor (ABCF), calculated as:

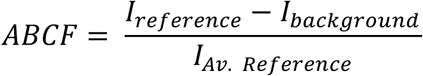

FRAP curves were normalized according to:

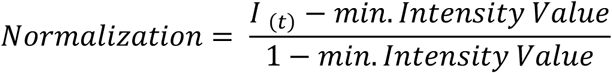

Values were averaged from 4 recordings for both early and late time points and the resulting FRAP curves ± standard deviation (std) were fitted for the early time points to a mono-exponential function:

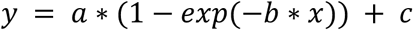

For the late time points, a bi-exponential function provided the best fitting:

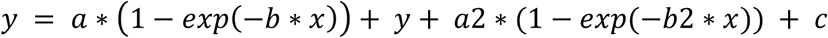

The mobile and immobile fractions were calculated using the parameters a and c derived from each fitting, according to the following equations:

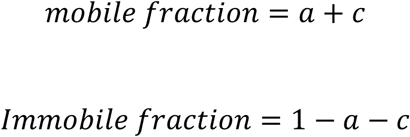

### NMR spectroscopy

NMR experiments were acquired on a 700MHz Bruker Avance III spectrometer equipped with a triple resonance TCI CryoProbe with Z-gradient and on a 800MHz Bruker Neo spectrometer equipped with a triple resonance TCI CryoProbe with Z-gradient. Experiments were performed at 278 K and 298 K. Chemical shifts of the NTE were assigned using a combination of two-dimensional TOCSY and NOESY NMR spectra and transferred to ^1^H-^15^N and ^1^H-^13^C HSQC to obtain N, Cα and Cβ chemical shifts. The hydrodynamic radius of the NTE was determined using pulse field gradient experiments with DSS used as internal reference. Spectra were processed using TopSpin 4.0.6 (Bruker) or NMRPipe 10.6 ^43^ and analyzed using SPARKY (Goddard TD & Kneller DG (2008) SPARKY 3. University of California, San Francisco).

Assigned chemical shifts were used to predict secondary structures in the NTE using TALOS+^28^.

### Molecular dynamics simulations

The starting structure of the N-terminal fragment of N^SARS-CoV-2^ was built with Flexible Meccano ^44^. Subsequently the peptide was equilibrated with 50,000 steps of energy minimization. To further equilibrate the system, 100 ps each of volume (NVT) and pressure (NPT) equilibration were performed. The MD simulations were carried out in GROMACS (version 2018.3) using the AMBER99SB-ILDN force field and the TIP3P water model at a temperature of 300 K, 1 bar of pressure and with a coupling time (ζT) of 0.1 ps. The mixtures were solvated in water with 150 mM NaCl, ensuring overall charge neutrality. The particle mesh Ewald algorithm was used for calculation of the electrostatic term, with a radius of 16 Å for the grid-spacing and Fast Fourier Transform. The cut-off algorithm was applied for the non-coulombic potential with a radius of 10 Å. The LINCS algorithm was used to contain bonds and angles. MD simulations were performed during 100 ns in 2 fs steps and saving the coordinates of the system every 10 ps. Five repetitions were done for each simulation, using a different structure from the ensemble of conformations generated by Flexible Meccano.

The hydrodynamic radius was calculated with the software HullRad (Version 7) ^46^. To derive the final hydrodynamic radius, the average of the radius of the structures of the last 50 ns of each simulation was taken together with the standard deviation.

## Supplementary Material

Supplementary material includes an analysis of LLPS of the full-length N^SARS-CoV-2^ and ΔNTE^SARS-CoV-2^ described by phase diagram and fluorescence recovery after photobleaching (FRAP). In addition, we include NMR spectra of N ^SARS-CoV-2^ amide and methyl spectral regions acquired at different concentrations of polyU.

## Acknowledgements

MZ was supported by the European Research Council (ERC) under the EU Horizon 2020 research and innovation program (grant agreement No. 787679).

## References

1. Chen H, Cui Y, Han X, Hu W, Sun M, Zhang Y, Wang P-H, Song G, Chen W, Lou J (2020) Liquid–liquid phase separation by SARS-CoV-2 nucleocapsid protein and RNA. Cell Res. 30:1143–1145.

2. Cubuk J, Alston JJ, Incicco JJ, Singh S, Stuchell-Brereton MD, Ward MD, Zimmerman MI, Vithani N, Griffith D, Wagoner JA, et al. (2021) The SARS-CoV-2 nucleocapsid protein is dynamic, disordered, and phase separates with RNA. Nat. Commun. 12:1–17.

3. Iserman C, Roden CA, Boerneke MA, Sealfon RSG, McLaughlin GA, Jungreis I, Fritch EJ, Hou YJ, Ekena J, Weidmann CA, et al. (2020) Genomic RNA Elements Drive Phase Separation of the SARS-CoV-2 Nucleocapsid. Mol. Cell 80:1078-1091.e6.

4. Mu J, Xu J, Zhang L, Shu T, Wu D, Huang M, Ren Y, Li X, Geng Q, Xu Y, et al. (2020) SARS-CoV-2-encoded nucleocapsid protein acts as a viral suppressor of RNA interference in cells. Sci. China Life Sci. 63:1413–1416.

5. Perdikari TM, Murthy AC, Ryan VH, Watters S, Naik MT, Fawzi NL (2020) SARS-CoV-2 nucleocapsid protein phase-separates with RNA and with human hnRNPs. EMBO J. 39:e106478–e106478.

6. Savastano A, Ibáñez de Opakua A, Rankovic M, Zweckstetter M (2020) Nucleocapsid protein of SARS-CoV-2 phase separates into RNA-rich polymerase-containing condensates. Nat. Commun. 11:6041.

7. Burbelo PD, Riedo FX, Morishima C, Rawlings S, Smith D, Das S, Strich JR, Chertow DS, Davey RT, Cohen JI (2020) Sensitivity in Detection of Antibodies to Nucleocapsid and Spike Proteins of Severe Acute Respiratory Syndrome Coronavirus 2 in Patients With Coronavirus Disease 2019. J. Infect. Dis. 222:206–213.

8. Dangi T, Class J, Palacio N, Richner JM, Penaloza MacMaster P (2021) Combining spike-and nucleocapsid-based vaccines improves distal control of SARS-CoV-2. Cell Rep. 36.

9. Lo YS, Lin SY, Wang SM, Wang CT, Chiu YL, Huang TH, Hou MH (2013) Oligomerization of the carboxyl terminal domain of the human coronavirus 229E nucleocapsid protein. FEBS Lett. 587:120–127.

10. Nikolakaki E, Giannakouros T (2020) SR/RS Motifs as Critical Determinants of Coronavirus Life Cycle. Front. Mol. Biosci. 7:219–219.

11. Bessa LM, Guseva S, Camacho-Zarco AR, Salvi N, Maurin D, Perez LM, Botova M, Malki A, Nanao M, Jensen MR, et al. (2022) The intrinsically disordered SARS-CoV-2 nucleoprotein in dynamic complex with its viral partner nsp3a. Science Advances 8:eabm4034.

12. Filho HVR, Jara GE, Batista FAH, Schleder GR, Tonoli CC, Soprano AS, Guimarães SL, Borges AC, Cassago A, Bajgelman MC, et al. (2021) Structural dynamics of SARS-CoV-2 nucleocapsid protein induced by RNA binding. bioRxiv:2021.08.27.457964-2021.08.27.457964.

13. Jack A, Ferro LS, Trnka MJ, Wehri E, Nadgir A, Nguyenla X, Fox D, Costa K, Stanley S, Schaletzky J, et al. (2021) SARS-CoV-2 nucleocapsid protein forms condensates with viral genomic RNA. PLOS Biol. 19:e3001425–e3001425.

14. Chang C-K, Hsu Y-L, Chang Y-H, Chao F-A, Wu M-C, Huang Y-S, Hu C-K, Huang T-H (2009) Multiple Nucleic Acid Binding Sites and Intrinsic Disorder of Severe Acute Respiratory Syndrome Coronavirus Nucleocapsid Protein: Implications for Ribonucleocapsid Protein Packaging. J. Virol. 83:2255–2264.

15. Dinesh DC, Chalupska D, Silhan J, Koutna E, Nencka R, Veverka V, Boura E (2020) Structural basis of RNA recognition by the SARS-CoV-2 nucleocapsid phosphoprotein. PLOS Pathog. 16:e1009100–e1009100.

16. Chang CK, Hou MH, Chang CF, Hsiao CD, Huang TH (2014) The SARS coronavirus nucleocapsid protein – Forms and functions. Antiviral Res. 103:39–50.

17. Wang J, Shi C, Xu Q, Yin H (2021) SARS-CoV-2 nucleocapsid protein undergoes liquid– liquid phase separation into stress granules through its N-terminal intrinsically disordered region. Cell Discov. 7:1–5.

18. Sprunger ML, Jackrel ME (2021) Prion-Like Proteins in Phase Separation and Their Link to Disease. Biomolecules 11:1014.

19. Alberti S, Halfmann R, King O, Kapila A, Lindquist S (2009) A Systematic Survey Identifies Prions and Illuminates Sequence Features of Prionogenic Proteins. Cell 137:146–158.

20. Lancaster AK, Nutter-Upham A, Lindquist S, King OD (2014) PLAAC: a web and command-line application to identify proteins with prion-like amino acid composition. Bioinformatics 30:2501–2502.

21. Tetz G, Tetz V (2018) Prion-like Domains in Eukaryotic Viruses. Sci. Rep. 8:1–10.

22. Syed AM, Taha TY, Tabata T, Chen IP, Ciling A, Khalid MM, Sreekumar B, Chen P-Y, Hayashi JM, Soczek KM, et al. (2021) Rapid assessment of SARS-CoV-2 evolved variants using virus-like particles. Science 0:eabl6184.

23. Davidson AD, Williamson MK, Lewis S, Shoemark D, Carroll MW, Heesom KJ, Zambon M, Ellis J, Lewis PA, Hiscox JA, et al. (2020) Characterisation of the transcriptome and proteome of SARS-CoV-2 reveals a cell passage induced in-frame deletion of the furin-like cleavage site from the spike glycoprotein. Genome Med. 12:1–15.

24. Bouhaddou M, Memon D, Meyer B, White KM, Rezelj VV, Correa Marrero M, Polacco BJ, Melnyk JE, Ulferts S, Kaake RM, et al. (2020) The Global Phosphorylation Landscape of SARS-CoV-2 Infection. Cell 182:685-712.e19.

25. Carlson CR, Asfaha JB, Ghent CM, Howard CJ, Hartooni N, Safari M, Frankel AD, Morgan DO (2020) Phosphoregulation of Phase Separation by the SARS-CoV-2 N Protein Suggests a Biophysical Basis for its Dual Functions. Mol. Cell 80:1092-1103.e4.

26. Guseva S, Perez LM, Camacho-Zarco A, Bessa LM, Salvi N, Malki A, Maurin D, Blackledge M (2021) ^1^H, ^13^C and ^15^N Backbone chemical shift assignments of the n-terminal and central intrinsically disordered domains of SARS-CoV-2 nucleoprotein. Biomol. NMR Assign. 15:255– 260.

27. Schiavina M, Pontoriero L, Uversky VN, Felli IC, Pierattelli R (2021) The highly flexible disordered regions of the SARS-CoV-2 nucleocapsid N protein within the 1–248 residue construct: sequence-specific resonance assignments through NMR. Biomol. NMR Assign. 15:219–227.

28. Shen Y, Delaglio F, Cornilescu G, Bax A (2009) TALOS+: a hybrid method for predicting protein backbone torsion angles from NMR chemical shifts. J. Biomol. NMR 44:213–223.

29. Marsh JA, Forman-Kay JD (2010) Sequence Determinants of Compaction in Intrinsically Disordered Proteins. Biophys. J. 98:2383–2390.

30. Abyzov A, Blackledge M, Zweckstetter M (2022) Conformational Dynamics of Intrinsically Disordered Proteins Regulate Biomolecular Condensate Chemistry. Chem. Rev. [Internet]. Available from: https://doi.org/10.1021/acs.chemrev.1c00774

31. Cascarina SM, Ross ED (2020) A proposed role for the SARS-CoV-2 nucleocapsid protein in the formation and regulation of biomolecular condensates. The FASEB Journal 34:9832– 9842.

32. Luo L, Li Z, Zhao T, Ju X, Ma P, Jin B, Zhou Y, He S, Huang J, Xu X, et al. (2021) SARS-CoV-2 nucleocapsid protein phase separates with G3BPs to disassemble stress granules and facilitate viral production. Science Bulletin 66:1194–1204.

33. Tompa P, Csermely P (2004) The role of structural disorder in the function of RNA and protein chaperones. The FASEB Journal 18:1169–1175.

34. Boncella AE, Shattuck JE, Cascarina SM, Paul KR, Baer MH, Fomicheva A, Lamb AK, Ross ED (2020) Composition-based prediction and rational manipulation of prion-like domain recruitment to stress granules. PNAS 117:5826–5835.

35. Scialò C, De Cecco E, Manganotti P, Legname G (2019) Prion and Prion-Like Protein Strains: Deciphering the Molecular Basis of Heterogeneity in Neurodegeneration. Viruses 11:261.

36. Seim I, Roden CA, Gladfelter AS (2021) Role of spatial patterning of N-protein interactions in SARS-CoV-2 genome packaging. Biophysical Journal 120:2771–2784.

37. Guseva S, Milles S, Jensen MR, Salvi N, Kleman J-P, Maurin D, Ruigrok RWH, Blackledge M (2020) Measles virus nucleo- and phosphoproteins form liquid-like phase-separated compartments that promote nucleocapsid assembly. Science Advances 6:eaaz7095

38. Sherry L, Smith M, Davidson S, Jackson D (2014) The N Terminus of the Influenza B Virus Nucleoprotein Is Essential for Virus Viability, Nuclear Localization, and Optimal Transcription and Replication of the Viral Genome. Journal of Virology 88:12326–12338.

39. He R, Leeson A, Ballantine M, Andonov A, Baker L, Dobie F, Li Y, Bastien N, Feldmann H, Strocher U, et al. (2004) Characterization of protein–protein interactions between the nucleocapsid protein and membrane protein of the SARS coronavirus. Virus Research 105:121–125.

40. Forsythe HM, Rodriguez Galvan J, Yu Z, Pinckney S, Reardon P, Cooley RB, Zhu P, Rolland AD, Prell JS, Barbar E (2021) Multivalent binding of the partially disordered SARS-CoV-2 nucleocapsid phosphoprotein dimer to RNA. Biophys. J. 120:2890–2901.

41. Krainer G, Welsh TJ, Joseph JA, Espinosa JR, Wittmann S, de Csilléry E, Sridhar A, Toprakcioglu Z, Gudiškytė G, Czekalska MA, et al. (2021) Reentrant liquid condensate phase of proteins is stabilized by hydrophobic and non-ionic interactions. Nat Commun 12:1085.

42. Liu X, Verma A, Garcia G, Ramage H, Lucas A, Myers RL, Michaelson JJ, Coryell W, Kumar A, Charney AW, et al. (2021) Targeting the coronavirus nucleocapsid protein through GSK-3 inhibition. Proc. Natl. Acad. Sci. U. S. A. 118.

43. Delaglio F, Grzesiek S, Vuister GW, Zhu G, Pfeifer J, Bax A (1995) NMRPipe: A multidimensional spectral processing system based on UNIX pipes. J. Biomol. NMR 6:277– 293.

44. Ozenne V, Bauer F, Salmon L, Huang J-R, Jensen MR, Segard S, Bernadó P, Charavay C, Blackledge M (2012) Flexible-meccano: a tool for the generation of explicit ensemble descriptions of intrinsically disordered proteins and their associated experimental observables. Bioinforma. Oxf. Engl. 28:1463–1470.

45. Fleming PJ, Fleming KG (2018) HullRad: Fast Calculations of Folded and Disordered Protein and Nucleic Acid Hydrodynamic Properties. Biophys. J. 114:856–869.

